# Metabolic reprogramming and elevation of glutathione in chromophobe renal cell carcinomas

**DOI:** 10.1101/649046

**Authors:** Yi Xiao, Rosanna Clima, Jonas Felix Busch, Anja Rabien, Ergin Kilic, Sonia Villegas, Seval Türkmen, Bernd Timmermann, Marcella Attimonelli, Klaus Jung, David Meierhofer

## Abstract

Chromophobe renal cell carcinomas (chRCC) are derived from intercalated cells of the collecting duct system, and are thought to be the malignant counterpart of benign renal oncocytomas. Here, we report the characterization of nine chRCC with adjacent healthy kidney tissues by applying proteome-, transcriptome (TCGA)-, and metabolome profiling. Most strikingly, the reactive oxygen species scavenger glutathione was significantly elevated in chRCC, caused by down-regulated enzymes involved in glutathione degradation. Metabolic reprogramming including stalled gluconeogenesis, down-regulated fatty acid- and amino acid metabolism was identified, even though, the abundance of amino acids and “energy carrier” molecules were unchanged. A striking anti-correlation of the mitochondrial respiratory chain between the transcriptome and the proteome was discovered, the transcripts coding for the respiratory chain were up-, while corresponding proteins and enzymatic activities were down-regulated. Similar to renal oncocytomas, chRCC exhibited a significant increase in glutathione, but are distinguishable by distinct regulation of the respiratory chain.

## Introduction

Chromophobe renal cell carcinomas (chRCC) are derived from intercalated cells of the collecting duct and comprise approximately 5% of all renal cancers (Moch et al., 2016). ChRCC show either a relatively transparent cytoplasm, or in about 30% of the cases, eosinophilic patterns with mitochondrial accumulation (Thoenes et al., 1988). The cytoplasm of chRCC is filled with a high number of vesicles, frequently with ‘inner vesicles’. Furthermore, mitochondria contain abundant tubulovesicular cristae and a few circular cristae, which appear scattered, and seem to originate from outer mitochondrial membrane outpouchings (Krishnan and Truong, 2002; Moreno et al., 2005; Thoenes et al., 1988).

One of the most characteristic genetic features of chRCC is monosomy of chromosomes 1, 2, 6, 10, 13, 17, and often 21 (Brunelli et al., 2005; Casuscelli et al., 2017; Haake et al., 2016). The most commonly mutated genes in chRCC are *TP53* (32%), *PTEN* (20%), and gene fusions involving the *TERT* promoter (Davis et al., 2014; Haake et al., 2016). Mutations in these tumor suppressors combined with the deletion of one of their chromosomes lead to a complete loss of function. Further mutations were observed with a lower frequency in *MTOR, NRAS, TSC1*, and *TSC2* indicating that the genomic targeting of the mTOR pathway occurred in 23% of all chRCC (Davis et al., 2014). Hence, *TP53’s* anticancer functions in apoptosis, genomic stability, and in inhibition of angiogenesis and PTEN’s role in the intracellular signaling pathway PI3K/AKT/mTOR are both disrupted, and can thus be regarded as major driving events in chRCC tumorigenesis. Mutations in these genes and imbalanced chromosome duplication were further shown to be important for the metastatic progression of chRCC and were associated with worse survival (Casuscelli et al., 2017).

In addition, mutations of the mitochondrial DNA (mtDNA) have been observed in complex I (CI) genes of the respiratory chain with a frequency of 21% (>20% heteroplasmy rate) (Davis et al., 2014). A pathway analysis showed that cases with mutations in *MT-ND5* versus no mutations in this gene in chRCC (almost all from the eosinophilic type) lead to an enrichment of the Gene Ontology term “mitochondrion” (Davis et al., 2014; Ricketts et al., 2018).

ChRCC are thought to be the malignant counterpart of benign renal oncocytomas, since they display many similarities: both are considered to be derived from the intercalated cell of the collecting duct, show frequent mtDNA mutations, and the eosinophilic type has an increased amount of mitochondrial mass. Disruption of Golgi and autophagy/lysosome trafficking events were attributed to defective mitochondrial function in renal oncocytomas (Joshi et al., 2015). Recently, metabolome profiling in chRCC revealed an impairment of the gamma-glutamyl transferase 1 activity and significantly increased levels of reduced and oxidized glutathione in chRCC (Priolo et al., 2018). A similar rewiring of the glutathione metabolism was also identified in renal oncocytomas (Gopal et al., 2018; Kurschner et al., 2017). Integrated proteome- and metabolome profiling datasets of chRCC are sparse. To fill this gap, we report here the results of 9 chRCC cases compared with adjacent healthy kidney tissues, obtained by profiling the proteome and metabolome. In addition, our results were correlated with existing chRCC transcriptome data and compared to our previous omics data from renal oncocytomas (Kurschner et al., 2017) to identify distinguishing markers.

## Results

### Proteome- and Metabolome Profiling of ChRCC

To comprehensively understand the nature of chRCC, we employed two systematic omics approaches; mass spectrometry-based proteome and metabolome profiling in nine chRCC and adjacent healthy kidney tissues (Table S1, Figure 1a, 1b). Distal and proximal tubules are known to be metabolically distinct, but we want to emphasize that they colocalize in any given section of the kidney, which serves as a healthy control in our study.

**Figure 1.**
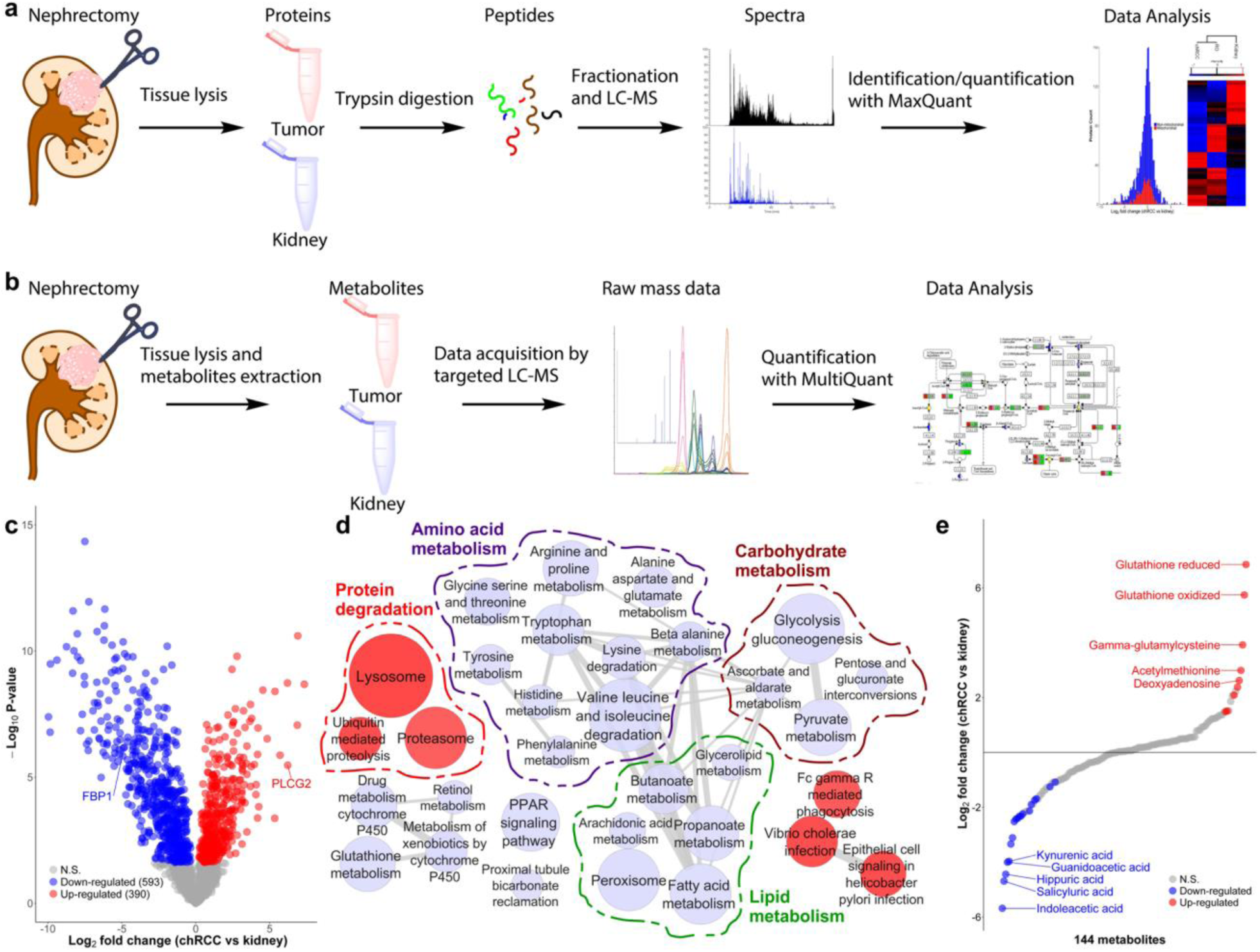
Proteome- and Metabolome Profiling of ChRCC. (a) The proteome workflow involves tissue lysis of the tumor and kidney samples, followed by a protein digestion with trypsin, peptide fractionation, analysis on the LC-MS and label-free quantification. (b) The metabolomics workflow involves tissue lysis of the tumor and kidney samples, metabolite extraction, followed by data acquisition by a targeted LC-MS approach and relative quantification of the peaks. (c) A volcano plot of log_2_ abundance ratios of chRCC versus kidney tissues (n =9) against the -log_10_ (p-value) of the proteome. Altogether 390 proteins were significantly up-regulated and are shown in red; 593 proteins were significantly down-regulated, as shown in blue. N.S., not significant. (d) Protein pathway analysis of chRCC versus kidney controls. Indicated are significantly (*p* ≤0.01 and FDR ≤ 0.15) enriched (red) and decreased (blue) KEGG pathways in chRCC. Grey lines connect overlapping pathways. Similar pathways are circled by a dotted line, such as decreased amino acid-, lipid-, carbohydrate metabolism, and increased protein degradation. (e) The distribution of fold changes of 144 metabolites in our cohort (n=9) of chRCC versus kidney tissues. Significantly (FDR <0.01) up-regulated and down-regulated metabolites are shown in red and blue, respectively. N.S., not significant.

A t-test with Benjamini-Hochberg (BH) correction (FDR < 0.05) for multiple testing was performed to identify significantly altered proteins between chRCC and controls. Altogether 983 significantly regulated proteins were identified, 390 proteins were significantly up- and 593 were down-regulated in chRCC (Figure 1c, Table S2). Among these significantly changed proteins, fructose-1,6-bisphosphatase (FBP1), which was shown to oppose clear cell RCC (ccRCC) progression (Li et al., 2014), decreased more than 16-fold in the tumor. Phospholipase C gamma 2 (PLCG2), whose high expression on transcriptome level had been considered as a potential biomarker for chRCC (Durinck et al., 2015), was over 59-fold increased in chRCC. Thus, our proteomic analysis could verify known molecular signatures of RCC.

A gene set enrichment analysis (GSEA) (Subramanian et al., 2005) was employed to assess whether an *a priori* defined set of proteins showed statistically significant, concordant differences between chRCC and healthy kidney tissues. GSEA revealed several significantly (*p* ≤0.01 and FDR ≤ 0.05) regulated Kyoto Encyclopedia of Genes and Genomes (KEGG) pathways in chRCC versus healthy tissues (Figure 1d, Table S3). The up-regulated pathways included the proteasome, ubiquitin-mediated proteolysis, the lysosome, and phagocytosis, in chRCC compared with normal tissue. In contrast, pathways involved in energy supply chains and nutrition homeostases, such as lipid metabolism, glutathione metabolism, the peroxisome, glycolysis and gluconeogenesis, and amino acid metabolism pathways were significantly down-regulated in chRCC (Figure 1 and Table S3).

A targeted metabolome profiling was performed and 144 metabolites were identified and relatively quantified (Figure 1e, Table S4). A statistical analysis using a two-sample t-test with the BH (FDR of 0.05) correction for multiple testing revealed 28 significantly (p-value <0.01) regulated metabolites (19 down-, 9 up-regulated) in chRCC versus kidney tissues (Figure 1e).

### Gluconeogenesis was Completely Stalled in ChRCC

The KEGG pathway “glycolysis and gluconeogenesis” was significantly reduced in chRCC (Figure 1d, Table S3). This pathway describes two opposing metabolic functions: the generation of pyruvate from glucose and vice versa. Thus, a more detailed view of each metabolic pathway showed that the abundance of all glycolytic enzymes were either unchanged or slightly increased, whereas all enzymes solely involved in gluconeogenesis were greatly reduced in chRCC, such as PC (124-fold), PCK1 (96-fold), PCK2 (996-fold), ALDOB (922-fold), and FBP1 (29-fold; Figure 2a). The two aldolase isoforms A and C, which were increased in chRCC, have a high affinity for fructose-1,6-bisphosphate (F-1,6-BP) to foster glycolysis, whereas the highly diminished isoform B has a low affinity for F-1,6-BP and hence converts the back-reaction from glyceraldehyde-3-phosphate to F-1,6-BP (Penhoet et al., 1966). Furthermore, the decreased PC, which is a main entry point for pyruvate to the TCA cycle, show that the tumor relies on lactate fermentation, also indicated by significantly increased LDHA (3-fold) levels in chRCC. These are therefore the most relevant metabolic changes observed in the tumor and can be regarded as a hallmark of chRCC. A comparison with gene expression data, extracted from The Cancer Genome Atlas (TCGA) data base (Davis et al., 2014), showed the same results, a significant depletion of transcripts coding for proteins involved in gluconeogenesis (Figure 2b-f).

**Figure 2.**
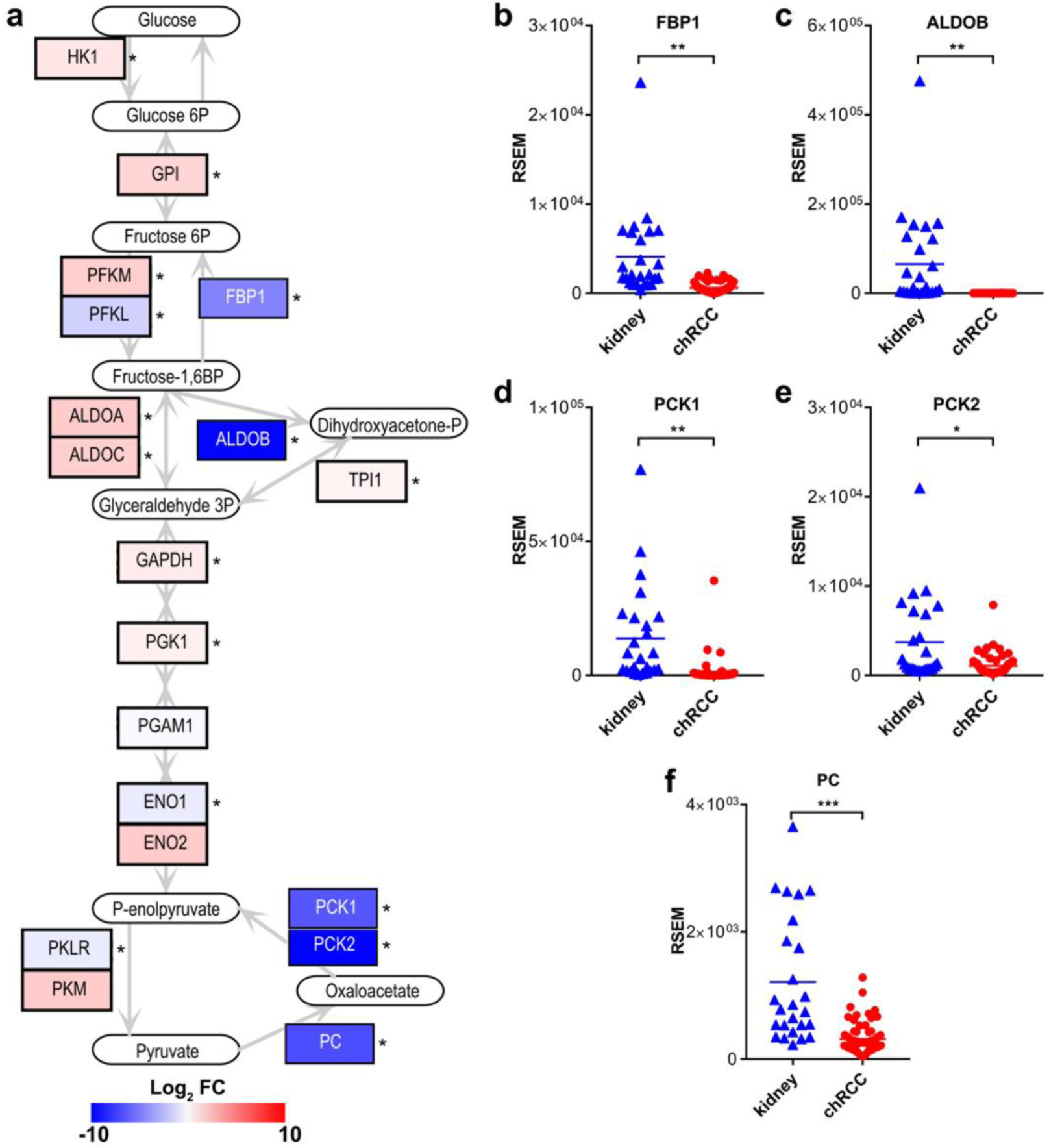
Stalled Gluconeogenetic Pathway in ChRCC. (a) Protein abundances of the metabolic pathways glycolysis and gluconeogenesis in chRCC. Log_2_ fold changes are displayed between chRCC and kidney tissues, up-regulated proteins are shown in red and down-regulated proteins in blue.* indicates significantly regulated proteins. (b-f) Transcript expression (TCGA) of gluconeogenic genes between chRCC (n= 66) and controls (n=25). (b) FBP1, fructose-1,6-bisphosphatase 1. (c) ALDOB, fructose-bisphosphate aldolase B. (d) PCK1, phosphoenolpyruvate carboxykinase, cytosolic. (e) PCK2, phosphoenolpyruvate carboxykinase, mitochondrial. (f) PC, pyruvate carboxylase. The data in (b-f) were calculated by RNA-Seq by Expectation Maximization (RSEM); *P < 0.05, **P < 0.01, ***P < 0.001 by a two-tailed student’s t-test.

### ChRCC Displays a Discrepancy Between the Abundance of Genes Coding for- and the Proteins Involved in the Respiratory Chain

By comparing the proteome profiles of chRCC and healthy tissues, we observed a general decrease in all subunits involved in the respiratory chain in chRCC, predominantly affecting CI subunits (Figure 3a). This is in clear contrast to previous reports based on transcript data from TCGA (Davis et al., 2014; Ricketts et al., 2018), which did not find a reduction of respiratory chain transcripts in this tumor (Figure 4a-b). We therefore compared our proteome to the transcriptome data from TCGA and found a high overall correlation between most proteins and transcripts, with a Pearson’s correlation coefficient of 0.671. However, the protein abundance of oxidative phosphorylation system (OXPHOS) subunits was negatively correlated with RNA expression in chRCC with a Pearson correlation coefficient of 0.169 (Figure 4c).

**Figure 3.**
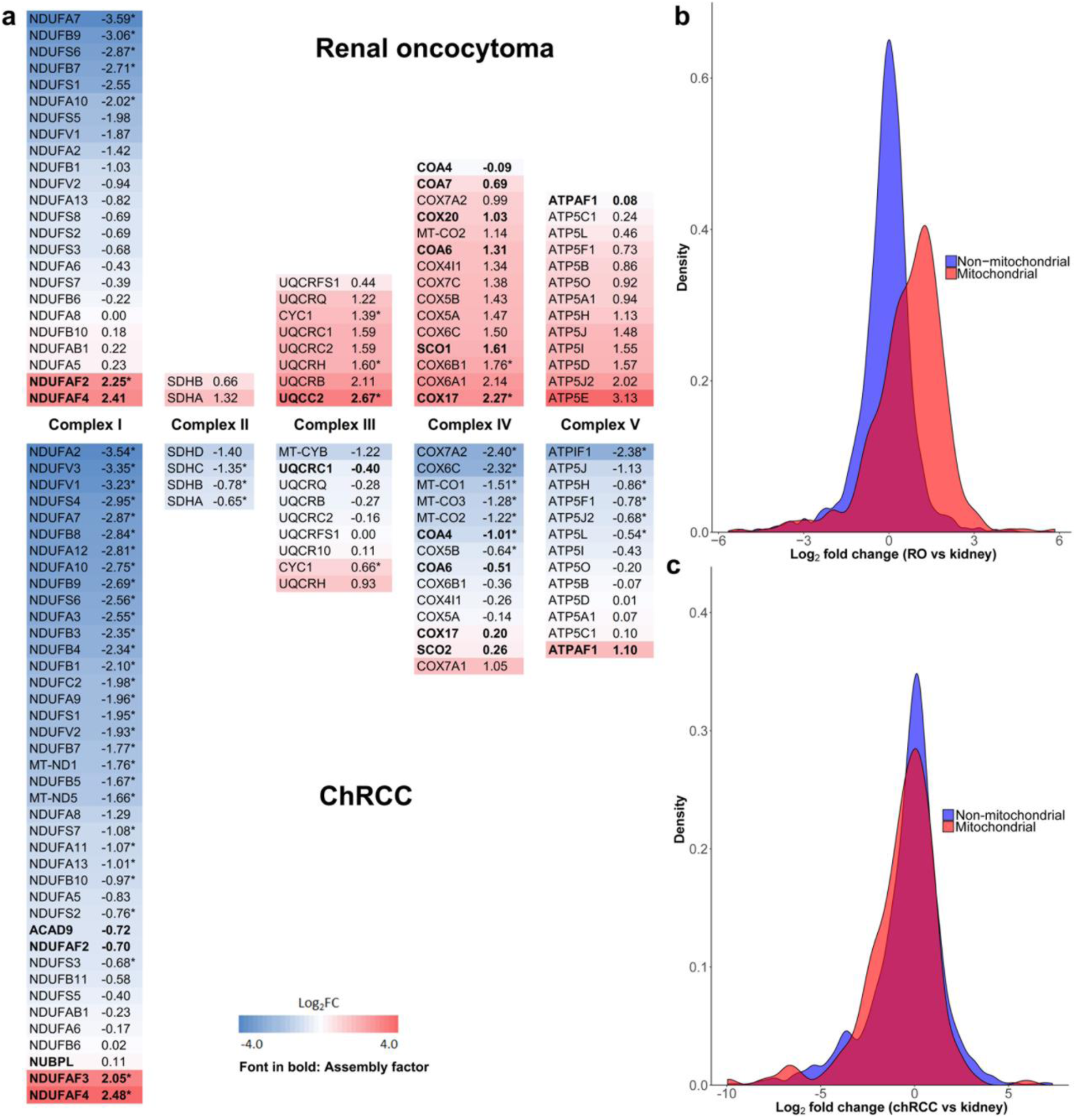
Distinct Regulation of the Respiratory Chain Complexes in Renal Oncocytomas and ChRCC. (a) Comparison of protein abundance ratios (log_2_) of all respiratory chain subunits between renal oncocytomas (top) and chRCC (bottom) samples versus healthy controls. Displayed are the four OXPHOS complexes and the F_o_-F_1_ ATPase, including all identified subunits and assembly factors and the corresponding log_2_ fold changes between chRCC and renal oncocytomas (Kurschner et al., 2017) versus kidney samples, respectively. The color gradient of the subunits reflects a low (blue) or high (red) abundance of this protein in the tumor. The abundances of CI subunits were most decreased in renal oncocytomas and chRCC, whereas all other respiratory chain complexes were increased in renal oncocytomas, but generally showing low abundance in chRCC assembly factors (in bold), not part of the final complex, were increased in all complexes in both tumor types. ^*^indicates significantly regulated proteins. (b) The density plot shows the density of mitochondrial and non-mitochondrial protein counts versus the log_2_ fold changes of proteins comparing renal oncocytomas versus kidney samples. (c) The density plot shows the density of mitochondrial and non-mitochondrial protein counts versus the log_2_ fold changes of proteins comparing chRCC versus kidney samples. Proteins located in mitochondria are indicated in red (Human Mito Carta, 1158 entries (Calvo et al., 2016)), whereas non-mitochondrial proteins are shown in blue.

**Figure 4.**
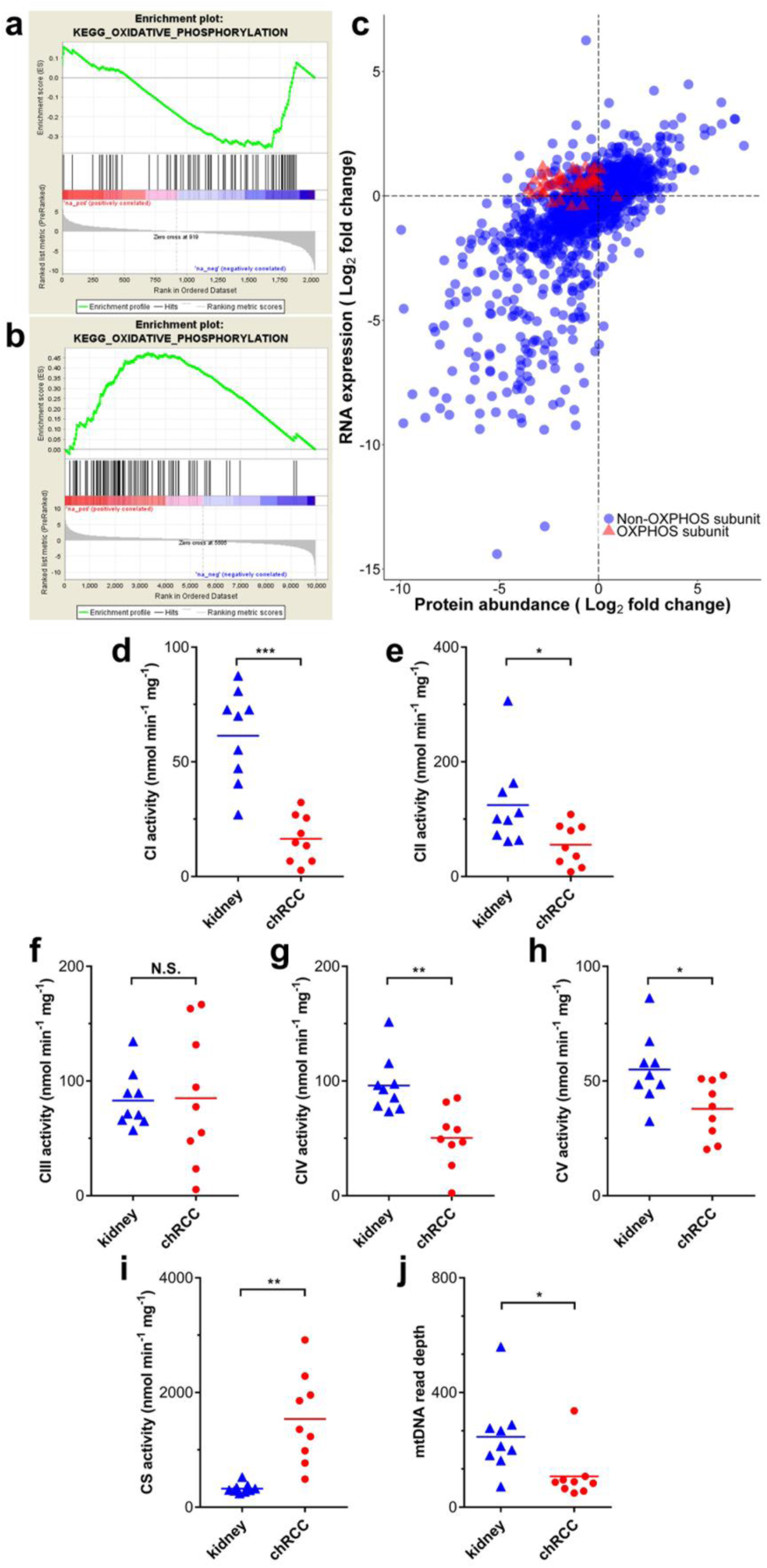
Comparison of the RNA Expression Versus the Protein Abundance Between ChRCC and Kidney Controls. (a) A pathway analysis (GSEA) shows the down-regulation of oxidative phosphorylation on the proteome level (this study). (b) A pathway analysis (GSEA) shows the up-regulation of the oxidative phosphorylation on the transcript level (TCGA data). (c) Indicated are genes/proteins involved in oxidative phosphorylation (red, triangles), which display the discrepancy between the expression of transcripts and protein abundance. RNA data were retrieved from TCGA. (d to i) Enzyme activities (nmol/min/mg protein, n =9) of the respiratory chain and TCA cycle in chRCC were compared with kidney controls. (d) Complex I (CI). (e) Complex II (CII). (f) Complex III (CIII). (g) Complex IV (CIV). (h) F_0_F_1_ ATPase (CV). (i) Citrate synthase (CS). (j) MtDNA abundances based on WES read depths (n=9). P-values in (d) to (j) are: *P < 0.05, **P < 0.01, and ***P < 0.001 by paired t-test. N.S., not significant.

In an additional step, the abundances of proteins involved in the OXPHOS system were compared with previously published data for benign renal oncocytomas (Kurschner et al., 2017), which are closely related to malignant chRCC. This revealed that only CI was decreased in both tumors, whereas all other respiratory chain complexes were only increased in renal oncocytomas (Figure 3a). An allocation analysis of mitochondrial versus non-mitochondrial proteins further highlighted the discrepancy between these tumor types showing that the mitochondrial protein mass was increased in renal oncocytomas (Figure 3b), whereas it was unchanged in chRCC (Figure 3c).

To validate the biological significance of the contradictory regulation between the transcriptome and proteome, the enzymatic activities of all respiratory chain complexes and citrate synthase (CS) were measured and the activities were compared for chRCC and the adjacent matching healthy tissues. This analysis revealed a significant reduction in the enzyme activities of CI, CII, CIV, and the F_0_F_1_ ATPase in chRCC, whereas CIII was unchanged and the activity of the TCA cycle enzyme CS was significantly increased (Figure 4d-4l).

Thus, the enzymatic activities of the respiratory chain reflect the abundance of protein rather than the expression of RNA transcripts. The TCA cycle, which is involved in many different metabolic pathways, seems to be regulated by a different mechanism since CS was significantly increased in chRCC. Nevertheless, the abundance of “energy carriers”, such as NAD^+^, NADH, NADP, ATP, and ADP was unchanged (Table S4).

### MtDNA Mutations and Copy Number Variations in chRCC

To examine whether mtDNA mutations are the main cause of the OXPHOS dysfunction in chRCC, as observed in renal oncocytomas (Kurschner et al., 2017; Mayr et al., 2008), we investigated the assembly of mitochondrial whole exome sequencing (WES) reads to identify somatic and germline mtDNA mutations in chRCC by pairwise comparison of healthy and tumor tissues. We found adequate coverage (>99.9%) and quality for a reliable mtDNA reconstruction and variant calling (Table S5). Although we identified five somatic non-synonymous events with a high disease score (>0.7), which are potentially pathogenic, it is not possible to infer a strong relationship with chRCC, considering that only one case showed a heteroplasmy of 60%, two below 30% and the remaining two just 5%. (Table S5).

To investigate, whether the presence or absence of mtDNA mutations (>50% heteroplasmy) leads to a different phenotype, we performed an analysis with the larger chRCC transcriptome cohort from TCGA (chRCC, n=66). A volcano plot showed no significantly changed transcripts between the mtDNA mutants and wild types (FDR <0.05, Figure S1), supporting our conclusion that mtDNA mutations do not play a primary role in chRCC progression.

By comparing the mtDNA reads between chRCC and controls, a significant and 3-fold decrease of the mtDNA content in chRCC was identified (Figure 4j). This has been previously observed for chRCC (Meierhofer et al., 2004) and might be directly correlated to the abundance decrease of OXPHOS subunits, as similar observations in ρ^0^ cells, entirely depleted of mtDNA, showed a severe reduction of all proteins involved in the respiratory chain (Aretz et al., 2016).

All WES reads were used to identify copy number variations (CNV) in the tumor. Monosomies of chromosomes 1, 2, 6, 10, 13, 17, and 21 were identified in all cases (Figure S1), conforming with previous reports (Brunelli et al., 2005; Casuscelli et al., 2017; Haake et al., 2016).

### ChRCC is Characterized by an Increased Level of Glutathione and Reduced Protein Abundance of Glutathione Degrading Enzymes

The top three increased metabolites in chRCC were all involved in glutathione metabolism, such as the ROS scavenger reduced glutathione (GSH, 115-fold), glutathione disulfide (GSSG, 54-fold), and γ-glutamylcysteine (15-fold; Figure 5a-c, Table S4). Case-specific GSH/GSSG ratios were found to be increased in all chRCC tissues, except for one case (Figure 5d). This indicates a reduced oxidative stress burden in chRCC, created by at least a hundred fold higher GSH and GSSG levels in chRCC compared with the kidney. In contrast to this, the glutathione metabolism pathway was significantly (p ≤ 0.005) reduced in chRCC. This was mainly due to decreased levels (average 42-fold) of glutathione degrading and conjugating enzymes, such as GSTM2, GGT5, GPX3, GGT1, GSTM3, GSTA1, and ANPEP as well as GPX3, which catalyzes the reduction of hydrogen peroxide. However, the enzymes involved in glutathione synthesis (GSS, GCLC), the glutathione peroxidases GPX1 and GPX4, the glutathione S-transferases GSTO1 and GSTP1, and mitochondrial glutathione reductase GSR remained unchanged. Therefore, the increase in reduced and oxidized glutathione in chRCC is due to significantly reduced abundance levels of proteins involved in glutathione degradation, rather than an increased GSH synthesis rate.

**Figure 5.**
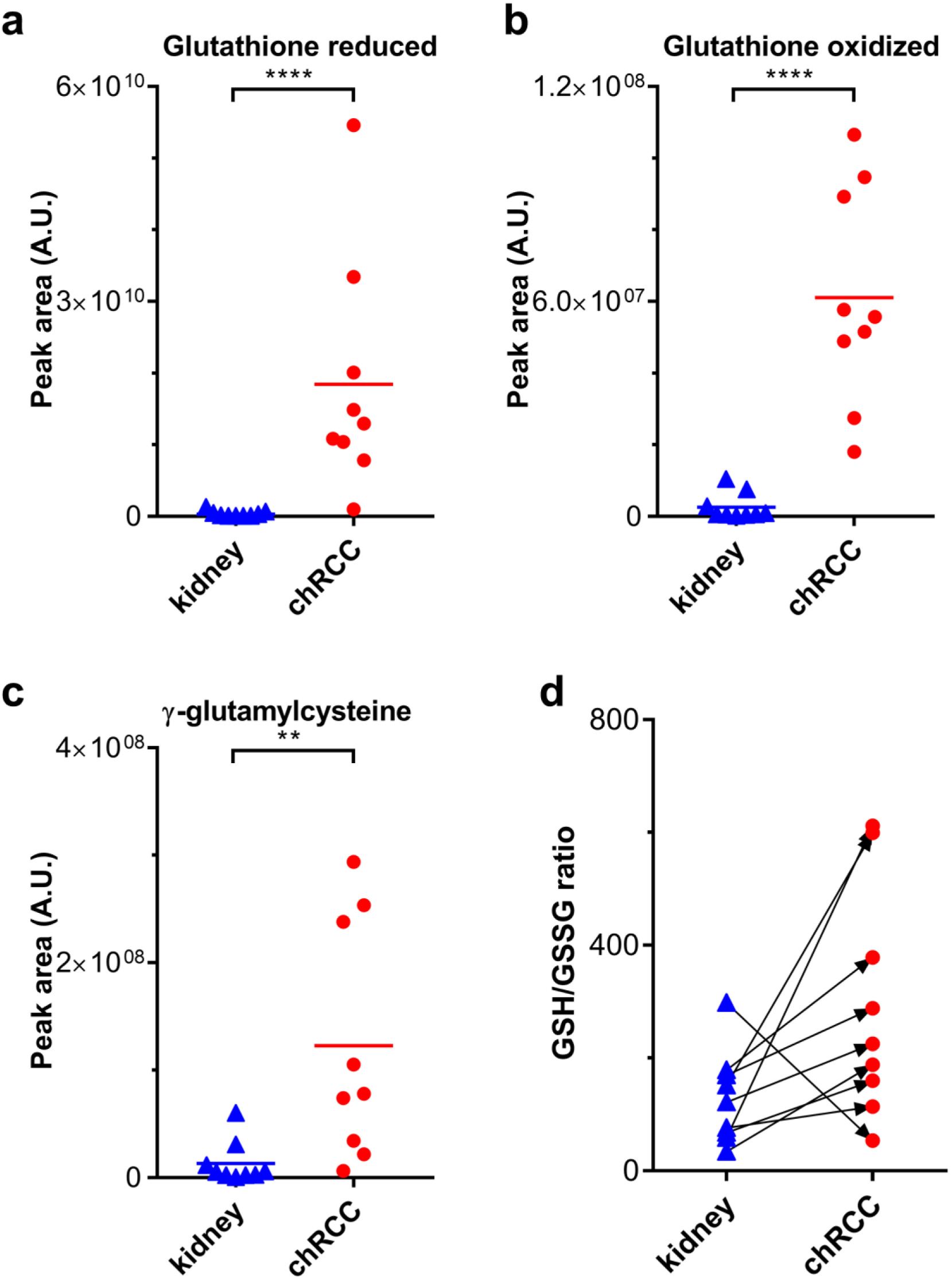
Profile of the Glutathione Metabolism in ChRCC. Relative abundances of metabolites involved in the glutathione metabolism are shown for normal kidney samples and chRCC. (a) Reduced glutathione. (b) Oxidized glutathione. (c) γ-glutamylcysteine. (d) GSH/GSSG ratio, calculated with the relative signal intensities. P-values in (*a*) to (*c*) are: **P < 0.001, ****P < 1e^−5^ by a two-tailed Student’s t-test. A.U., arbitary units.

In a pilot study, we further analyzed free and total GSH levels in plasma and urine of ccRCC, pRCC, chRCC, and renal oncocytoma specimens to elucidate whether GSH can be used as a non-invasive diagnostic marker. The results clearly indicated that free (Figure S2 a,c) and total (Figure S2 b,d) GSH cannot serve as a metabolic marker to distinguish between renal tumor types and healthy individuals in urine and plasma.

### ChRCC Features a Depletion of Amino Acid Intermediates and Pathways Involved in Amino Acid Metabolism

On the proteome level, ten distinct pathways involved in amino acid metabolism were significantly down-regulated in chRCC (Figure 1d, Table S3). In total, 95 out of 114 proteins involved in amino acid metabolism showed a decreased abundance (average 22-fold). Altogether 19 proteins even decreased in abundance by over 40-fold (BBOX1, HPD, PSAT1, ACY1, DDC, ALDH3A2, CHDH, MAOB, FTCD, EHHADH, DPYS, DMGDH, ALDH4A1, SHMT1, ABAT, ASS1, AGMAT, BHMT, and GATM), indicating a major metabolic change (Figure 6a). In addition, six pathways associated with fatty acid metabolism were significantly down-regulated (Figure 1d, Table S3), including fatty acid metabolism and the peroxisome, the main organelle for fatty acid oxidation. The abundances of amino acid intermediates were found to be among the most depleted metabolites in chRCC, matching the decreased protein abundances of amino acid pathways. These included guanidoacetic acid (intermediate of multiple amino acids, glycine, serine, threonine, arginine and proline, 16-fold), kynurenic acid (tryptophan intermediate, 16-fold), indoleacetic acid (tryptophan intermediate, 51-fold), ureidopropionic acid (beta-alanine intermediate, 6-fold), hippuric acid (glycine intermediate, 22-fold), salicyluric acid (glycine intermediate, 26-fold) and acetylglutamine (glutamine intermediate, 5-fold) (Figure 6b-6g, Table S4). Interestingly, the level of glycine was not significantly decreased, but all of the other amino acids were either unchanged or slightly, but not significantly increased. Selected amino acids and intermediates were mapped together with according enzymes abundances onto their pathways to indicate whether they were derived up- or down-stream (Figure S3).

**Figure 6.**
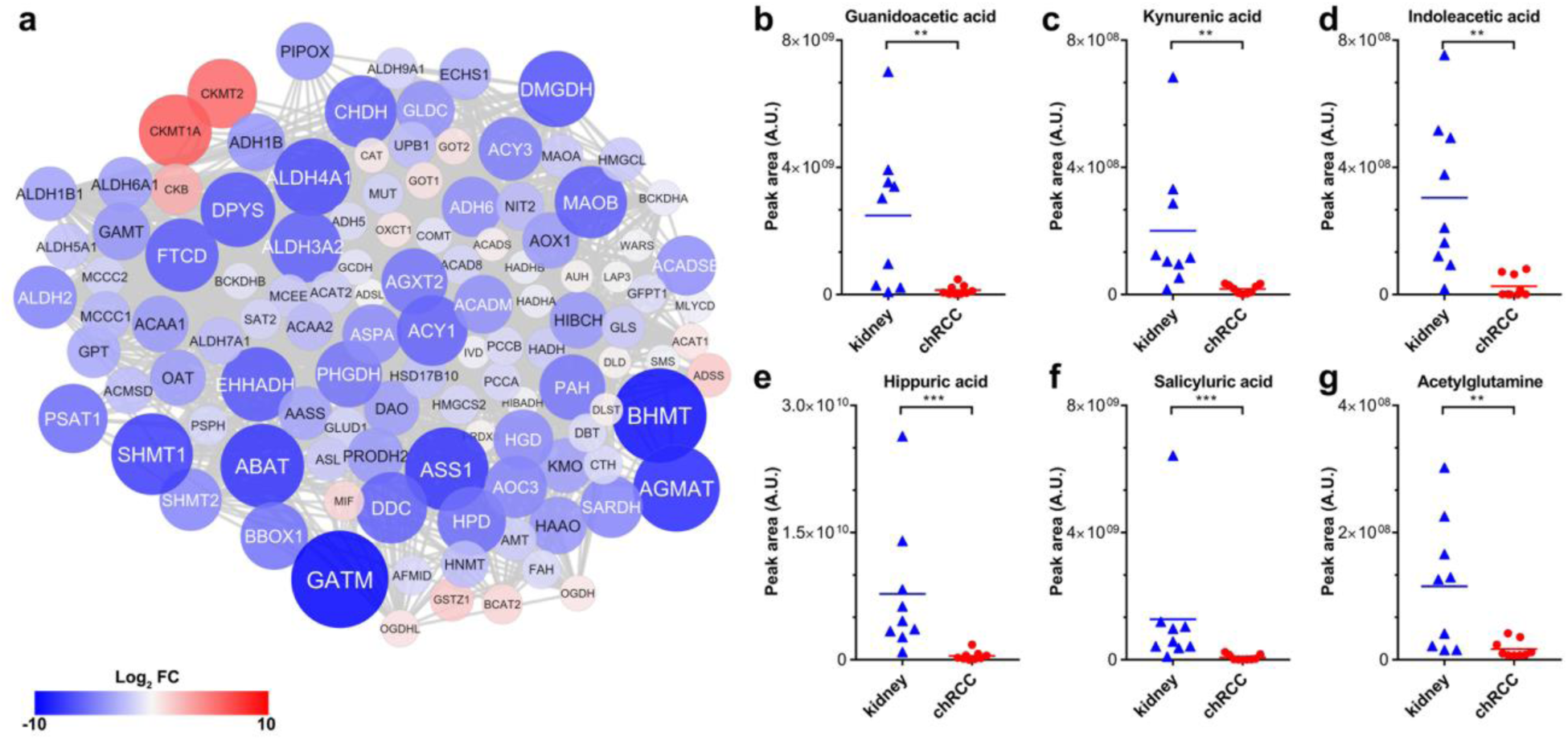
Regulation of Amino Acid Metabolism in ChRCC. (a) A protein-protein interaction network was created to elucidate the regulation of the entire amino acid metabolism in chRCC. The colors of the nodes correspond to the protein expression fold change comparing chRCC versus healthy kidney tissues; red indicates higher expression and blue lower expression in chRCC. The size of the nodes corresponds to the absolute protein expression fold-change. (b-g) The relative abundance of six selected amino acid intermediates are shown for kidney tissues and chRCC. (b) Guanidoacetic acid. (c) Kynurenic acid. (d) Indoleacetic acid. (e) Hippuric acid. (f) Salicyluric acid. (g) Acetylglutamine. P-values in (b) to (g) are: **P < 0.001, ***P < 5e^−4^ by two-tailed Student’s t-test. A.U., arbitary units.

### ChRCC Cells Feed on Extracellular Macromolecules Via Endocytosis

Pathways related to amino acid metabolism were significantly downregulated in chRCC (Figure 1d, Figure 6), whereas all detected amino acid levels were unchanged (Table S4). Thus, we hypothesized that chRCC cells could preferentially internalize and catabolize external macromolecules as a source of amino acids. Indeed, a pathway related to endocytosis (Fc gamma R mediated phagocytosis) and three protein degradation pathways (lysosome, ubiquitin-mediated proteolysis, and proteasome) were significantly up-regulated in chRCC (Figure 1d, Table S3). In particular, the two lysosome markers, LAMP1 and LAMP2, were up-regulated by 3 and 7-fold, respectively (Table S2). Activated lysosomes indicate that recycling of macromolecules and maintaining an acidic tumor microenvironment favors the viability and progression of chRCC. These metabolic chances indicate that extracellular biomass recruitment and protein degradation is the preferred way of gaining mass and nutrition in chRCC.

To further investigate, whether increased abundances of lysosomal proteins correlate with increased enzymatic activities in chRCC, we measured the activity of two lysosomal enzymes, hexosaminidase A and B. These enzymes are involved in the breakdown of gangliosides and were both found to be significantly increased in chRCC (Figure 7a-b).

**Figure 7.**
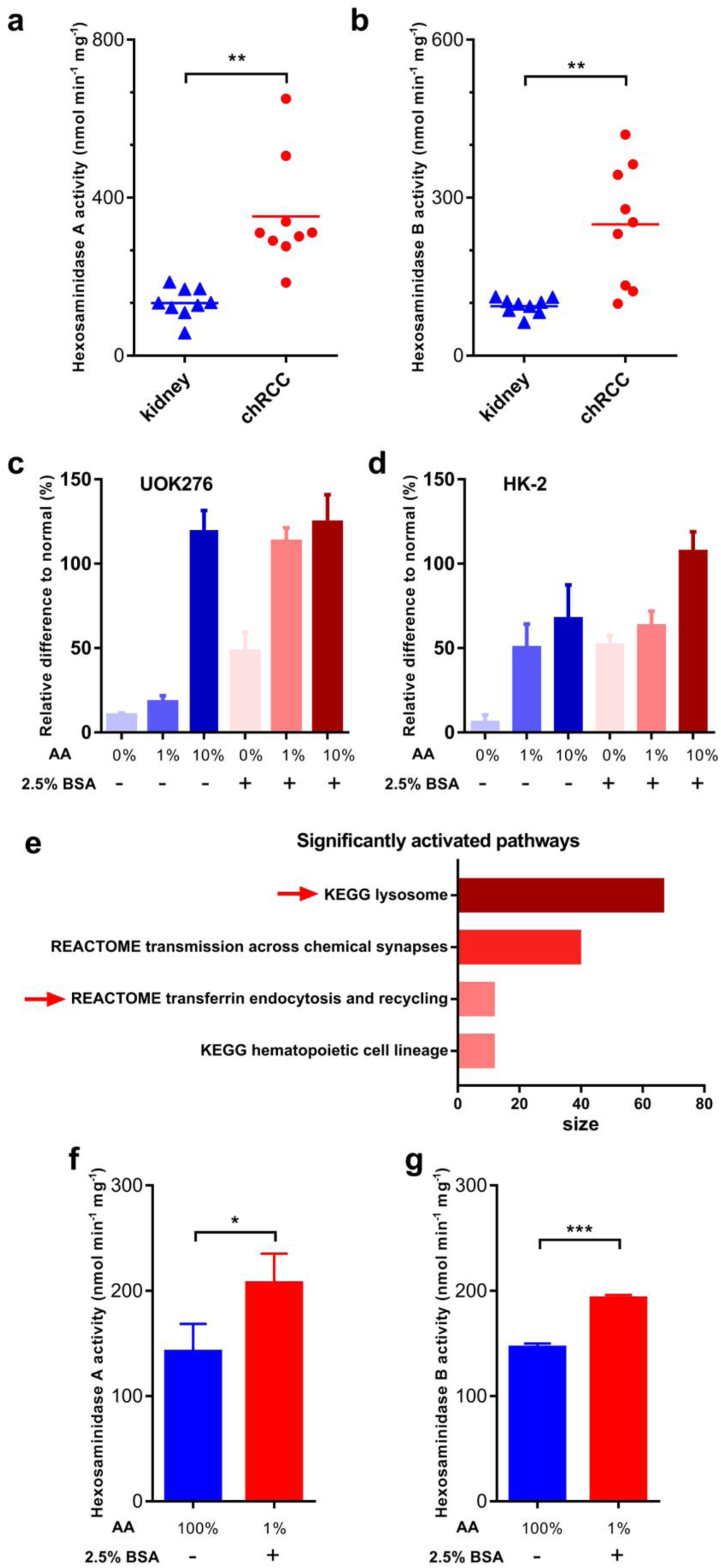
ChRCC Cells Activated Endocytosis when Amino Acids were Depleted. (a-b) Enzymatic activities (nmol/min/mg protein, n =9) of the hexosaminidase A (a) and B (b) in chRCC compared with kidney controls. (c-d) Relative cell numbers of UOK276 cells (c) and HK-2 cells (d) under different amino acid (AA) and bovine serum albumin (BSA) concentrations normalized to complete medium. (e) A pathway analysis (GSEA) of the proteome shows the up-regulation of the lysosome and endocytosis in UOK276 cells in 1% amino acid and 2.5% BSA versus a complete medium, enriched pathway cutoff, p < 0.01, FDR < 0.25. (f-g) Enzymatic activities (nmol/min/mg protein, n =3) of the hexosaminidase A (f) and B (g) in UOK276 cells in complete medium without supplemetation of BSA and 1% amino acids with 2.5% BSA. The data in (c-d) and (f-g) are expressed as means ± SD, p-values in (a-b) are: **P < 0.01, by paired t-test; p-values in (f-g) are: *P < 0.01, **P < 0.01, two-tailed Student’s t-test.

Next, we asked, whether the supplementation of a macromolecule indeed enhances the growth rate of chRCC cells and whether this is mirrored by the alteration of protein abundances in endocytosis pathways. Therefore, we evaluated the proliferation of the chRCC-derived cell line UOK276 (Yang et al., 2017) and normal kidney cells (HK-2) under different amino acid and bovine serum albumin (BSA) concentrations. Overall, both UOK276 and HK-2 cells grew significantly faster under all amino acid concentrations when supplemented with 2.5% BSA, only the UOK276 cells grew equally well when feed with 10% amino acids (Figure 7c-d). UOK276 cells proliferated normally under a low amino acid concentration of 1% and the addition of 2.5% BSA, whereas HK-2 cells significantly reduced the growth rate. This result suggests that UOK276 cells have a higher potential to utilize BSA to compensate the amino acid depletion. Proteome profiling between UOK276 cells supplemented with 1% amino acids and with the addition of 2.5% BSA versus a complete medium revealed a significantly increase of the pathways endocytosis (transferrin endocytosis and recycling, Reactome) and the lysosome (KEGG) (Figure 7e, Table S6). An enzymatic activity measurement of the lysosomal hexosaminidases A and B of UOK276 cells between the before mentioned conditions further confirmed that external macromolecules triggered the internalization and the degradation pathways to break down macromolecules enzymatically via endocytosis (Figure 7f-g), which indicates the adaption of chRCC cells to nutrient-poor conditions. Hence, the experiments performed in the chRCC derived UOK276 cell line validated the observations gained for the chRCC tissues.

## Discussion

The first description of chRCC as a new kidney cancer entity by Thoenes et al. (Thoenes et al., 1985) was over 30 years ago. Recently, a few studies, comprising roughly a hundred cases (Davis et al., 2014; Ricketts et al., 2018), were undertaken to understand the genetic cause of chRCC. In contrast, proteome and metabolome profiling data from this malignant tumor are still sparse (Priolo et al., 2018; Schaeffeler et al., 2018; Valera et al., 2010). To fill this gap, this study employs and integrates proteomics, transcriptomics (from TCGA), and metabolomic approaches as well as mitochondrial WES to gain insights into the mtDNA mutation landscape and to elucidate metabolic changes between chRCC and adjacent kidney tissues. In addition, these results were compared with renal oncocytomas data from our previous study (Kurschner et al., 2017) to identify molecular differences between these highly similar tumor entities at the proteome- and metabolome level.

The most strikingly increased set of metabolites in chRCC were those involved in glutathione metabolism (GSH, GSSG, γ-glutamyl-cysteine, Figure 2). This is similar to those previously identified in chRCC (Priolo et al., 2018), renal oncocytomas (Gopal et al., 2018; Kurschner et al., 2017), and in ccRCC (Hakimi et al., 2016; Wettersten et al., 2015). Since GSH is an important ROS scavenger (Circu and Aw, 2008) the increased GSH levels and GSH/GSSG ratios in chRCC can thus be considered as the main strategy for the tumor to overcome ROS stress originating from a dysregulated respiratory chain. Similar to renal oncocytomas, enzymes involved in glutathione synthesis were unchanged, but enzymes involved in glutathione degradation were significantly reduced in chRCC.

The second hallmark in chRCC was the substantial reprogramming of the central metabolic pathways. Glycolysis and gluconeogenesis share most of the enzymes, which can run in both directions, but only the abundance of gluconeogenesis specific enzymes were significantly and up to thousand-fold reduced in chRCC (Figure 2). The kidney is the only organ, besides the liver, which can deliver anabolic glucose for the organism, but a net flux can be only generated for one direction to avoid a futile cycle as this would violate thermodynamics. All kidney tumors seem to abandon this energy consuming pathway, since similar observations were also reported in renal oncocytomas (Kurschner et al., 2017) and at the transcript level for pRCC (Cancer Genome Atlas Research et al., 2016) and ccRCC (Li et al., 2014).

The down-regulation of the fructose-1,6-bisphosphatase 1 gene (*FBP1*), a key player in gluconeogenesis, was previously linked to ccRCC progression and was shown to inhibit nuclear hypoxia-inducible factor function (Li et al., 2014). This further stimulates the metabolic switch by up-regulating the glycolytic target genes upon its loss in the tumor. This effect of increasing glycolytic enzymes in chRCC was also seen in our proteome data (Figure 3). Of equal importance is the uptake of glutamine and glucose to sustain tumor growth, which was shown to be significantly lower in chRCC compared with ccRCC and pRCC (Nakajima et al., 2017). This could be the consequence of a low micro-vessel density observed in chRCC compared with ccRCC tumors (Jinzaki et al., 2000). Hence, chRCC seem to be poorly supported by classical nutrition supply chains.

Contrary to the observed abundance decrease of gluconeogenic proteins in chRCC, the over-expression of gluconeogenic genes and proteins can be frequently found in other tumor species. Specifically, increased *ALDOB* expression was found to favor cancer cell proliferation and metastasis in multiple cancers, such as colon cancer (Bu et al., 2018), rectal cancer (Tian et al., 2017), and colorectal adenocarcinoma (Li et al., 2017). This might also be one reason for the low metastatic potential (Volpe et al., 2012) and the high survival rate of chRCC patients (Amin et al., 2002; Przybycin et al., 2011).

An increase in glycolytic activity frequently goes hand-in-hand with reduced oxidative phosphorylation, even under aerobic conditions, known as the Warburg effect. This phenomenon was also observed in our chRCC cohort. The abundance of all oxidative phosphorylation complexes and the F_0_F_1_-ATPase and most of their enzymatic activities were significantly reduced (Figure 4). Previously, we found a significant reduction in all mitochondrial enzyme activity as well as in the mtDNA content in ccRCC and pRCC (Meierhofer et al., 2004). At this specific point, we see a clear difference to benign renal oncocytomas, where only CI subunits and its enzyme activity were significantly reduced, while all other complexes and enzyme activities were significantly increased (Kurschner et al., 2017).

How closely transcript levels correspond to protein abundances is a fundamental question. A discrepancy between increased transcript expression (Figure 4) and reduced protein abundances observed for respiratory chain genes and proteins has not only been observed in our chRCC cohort but also previously in renal oncocytomas (Kurschner et al., 2017). This anti-correlation indicates that different levels of molecular information are necessary to evaluate and understand a pathomechanism. Thus, the enzymatic activities of the respiratory chain in chRCC and also in renal oncocytomas (Mayr et al., 2008) matched to protein abundances rather than to gene expression.

The mitochondrial genomes in chRCC and thyroid cancers have been shown to have a particularly enriched mutation load (Davis et al., 2014; Grandhi et al., 2017). Although we identified diverse potentially pathogenic mutations with mostly low heteroplasmy loads, we conclude that they most likely do not play a primary role in the pathogenesis and progression of chRCC, as the presence or absence of mtDNA mutations did not alter the abundance of OXPHOS subunits. This phenomenon has been shown before in chRCC and alternative roles, such as a compensatory role for the loss of complex I function or selective pressures operating to promote alternative pathways have been suggested (Davis et al., 2014).

Beside the dramatic reduction in enzymes involved in gluconeogenesis and the respiratory chain, a decrease in the proteins of the fatty acid and amino acid synthesis pathways was identified in chRCC (Figure 1d). Interestingly, the actual level of amino acids and metabolites, which can be regarded as essential “energy carriers”, such as FAD, NADH, ADP, cyclic AMP, and NAD^+^ were indeed unchanged (Table S4), indicating a sufficient amount of building blocks and nutrients in chRCC. Only amino acid intermediates were decreased in chRCC, reflecting a lower activity of pathways involved in amino acid metabolism (Figure 6b-6g). In contrast, increased amino acid metabolism was frequently found in other cancer types to promote proliferation and metastasis, such as glycine and serine metabolism in breast cancer (Jain et al., 2012).

Based on these findings, we hypothesize that chRCC acquire nutrients preferentially in a different way: The lysosome and the proteasome, which are involved in the cell recycling machinery by contributing and delivering new biomass via endocytosis, and autophagy were significantly enriched (Figure 1d). Indeed, the chRCC derived cell line UOK276 could utilize macromolecules better under nutrient-poor conditions (Figure 7) and supplementation of BSA triggered a significant increase of the lysosome. The reduced abundance of pathways involved in amino acid metabolism paralleled with an increased abundance of the proteasome, ubiquitin-mediated proteolysis, and the lysosome suggests a main adaptive mechanism to production, recycling, and energy metabolism in chRCC, such as facilitating the maintenance of proteostasis (Zhang and Manning, 2015) or responding to stress (Livneh et al., 2016).

Furthermore, endocytosis was reported to suppress cancer cell blebbing and invasion by increasing the cell volume and membrane tension and thus decrease the likelihood of mechanical invasion of the surrounding tissue (Holst et al., 2017). This further matches the low rate of chRCC patients (1.3%), who develop distant metastases (Volpe et al., 2012) and correlates with our finding of enriched endocytosis. The modulation of the microenvironment by the acidity of vascular ATPases, which also increased in chRCC, to acidify and hydrolyze macromolecules to fuel biomass production (Davidson and Vander Heiden, 2017), and for the extracellular space (Dettmer et al., 2006; Repnik et al., 2013) might be another advantage for chRCC survival and viability.

A histopathological differentiation of benign renal oncocytomas from malignant chRCC is still difficult, even using multiple immunohistochemistry markers (Ng et al., 2016). There are also case reports on rare hybrid tumors (Noguchi et al., 1995), including oncocytic content (Pote et al., 2013) and the rare genetic disorder Brit-Hogg-Dube (BHD) syndrome (Vera-Badillo et al., 2012), where the coexistence of renal oncocytomas and chRCC was shown. The defective mitochondriogenesis was named as a possible source of microvesicles in chromophobe renal cell carcinomas. But microvesicles are also not diagnostic for chRCC, as they can also be found in renal oncocytomas and eosinophilic variants of ccRCC (Tickoo et al., 2000). Consequently, the abundance signature of OXPHOS complexes could be used to unambiguously distinguish these two tumor species, since these pathways were regulated in opposing directions (Figure 3). On the metabolite level, amino acids could be used to distinguish both tumor entities; a decrease was found in renal oncocytomas (Kurschner et al., 2017) and no change was seen in chRCC. In summary, ChRCC are characterized by substantially increased levels of the ROS scavenger glutathione with decreased gluconeogenesis and respiratory chain activity. We therefore hypothesize that increased protein degradation, and lysosomal and proteasomal activities are necessary for chRCC to acquire nutrition from the extracellular matrix in order to compensate and counteract the lower glucose uptake, lower amino acid synthesis rates, and respiratory chain activities. Moreover, the abundance of proteins involved in the respiratory chain could be used as a biomarker to distinguish between benign oncocytomas and malignant chRCC.

## Acknowledgements

This work is part of the doctoral dissertation of Y.X. Our work is supported by the Max Planck Society and the China Scholarship Council to Y.X. and the Foundation for Urologic Research (A.R. and K.J.). We are thankful to Prof. Marston Linehan, Center for Cancer Research, Bethesda, for providing the chRCC cell line UOK276.

## Author Contributions

Y.X. performed omics profiling, analyzed the data and prepared figures; J.F.B. and A.R. recruited chRCC cases, E.K. and S.V. validated histological samples, B. T. acquired WES data, R.C. and M.A. analyzed WES data; S.T. performed karyograms; Y.X and D.M. wrote the manuscript; D.M. conceived and directed the project, A.R. and K.J. reviewed the manuscript.

## Declaration of Interests

The authors declare no competing interests.

## Methods

### Tissue Dissection and Verification of ChRCC

Nine pairs of chRCC and their non-malignant counterparts derived from nephrectomies were collected in liquid nitrogen immediately after surgery and preserved at −80°C. The clinical characteristics of the tumors are reported in Table S1. From the collected tissue samples, histologic sections were stained with hematoxylin and eosin. The diagnosis of chRCC and the corresponding matched tumor-free kidney tissue was done according to the WHO classification criteria. Only cases with a clear diagnosis of chRCC were considered for the study. The study was approved by the institutional Ethics Committee (no. EA1/134/12) and was carried out in accordance with the Declaration of Helsinki. All participants gave informed consent.

### Cell Culture Conditions for the Proliferation Assay

The recently established and chRCC-derived cell line UOK276 (Yang et al., 2017) and the human kidney (HK-2, cortex/proximal tubule, ATCC CRL-2190) cell line were cultivated in Dulbecco’s modified Eagle medium (DMEM, Life Technologies, New York, NY) containing 4.5 g/L glucose, supplemented with 10% fetal bovine serum (FBS, Silantes, Munich, Germany) and 1% penicillin–streptomycin–neomycin (Invitrogen, Carlsbad, CA) at 37 °C in a humidified atmosphere of 5% CO_2_.

For the proliferation assays, UOK276 and HK-2 cells were seeded in 96-well plates at a density of 2×10^4^ cells per well in complete medium. After 12 hours, cells were briefly rinsed and cultured in minimum essential medium (MEM) supplemented with 10% dialyzed FBS (Thermo Fisher, Germany), amino acid (0, 1, and 10% versus MEM), and BSA (0 and 2.5% total) concentrations. The number of cells was determined after 4 days.

### Tissue Sample Preparation for Proteomics

The tissue samples for proteomics were processed with iST 96X kits following the manufacturer’s protocol (iST sample preparation kit 96X, PreOmics, Martinsried, Germany). Briefly, tissues were homogenized under denaturing conditions with a FastPrep (three times for 60 s, 6.5 m x s^−1^), followed by boiling at 95°C for 10 min. The lysates containing 40 µg protein were then digested by trypsin and Lys-C protease mixture at 37 *°*C overnight. Subsequently, the peptides were purified with the cartridge and each sample was further separated using three fractions according to (Kulak et al., 2014). A total of 10 µg of each fraction were analyzed by LC-MS for proteome profiling. All fractions were allocated to the corresponding replicate and analyzed jointly by the software tool MaxQuant (Cox and Mann, 2008).

Proteomic profiling of the UOK276 cells was done as reported previously (Kurschner et al., 2017) in biological triplicates.

### LC-MS Instrument Settings for Shotgun Proteome Profiling and Data Analysis

LC-MS/MS was carried out by nanoflow reverse phase liquid chromatography (Dionex Ultimate 3000, Thermo Scientific, Waltham, MA) coupled online to a Q-Exactive HF Orbitrap mass spectrometer (Thermo Scientific, Waltham, MA). Briefly, the LC separation was performed using a PicoFrit analytical column (75 μm ID × 55 cm long, 15 µm Tip ID (New Objectives, Woburn, MA) in-house packed with 3-µm C18 resin (Reprosil-AQ Pur, Dr. Maisch, Ammerbuch-Entringen, Germany). Peptides were eluted using a gradient from 3.8 to 40% solvent B in solvent A over 120 min at 266 nL per minute flow rate. Solvent A was 0.1 % formic acid and solvent B was 79.9% acetonitrile, 20% H_2_O, 0.1% formic acid. Nanoelectrospray was generated by applying 3.5 kV. A cycle of one full Fourier transformation scan mass spectrum (300-1750 m/z, resolution of 60,000 at m/z 200, AGC target 1e^6^) was followed by 16 data-dependent MS/MS scans (resolution of 30,000, AGC target 5e^5^) with a normalized collision energy of 27 eV. In order to avoid repeated sequencing of the same peptides, a dynamic exclusion window of 30 sec was used. In addition, only peptide charge states between two to eight were sequenced.

Raw MS data were processed with MaxQuant software (v1.6.0.1) (Cox and Mann, 2008) and searched against the human proteome database UniProtKB with 70,941 entries, released in 01/2017. Parameters of MaxQuant database searching were: A false discovery rate (FDR) of 0.01 for proteins and peptides, a minimum peptide length of 7 amino acids, a mass tolerance of 4.5 ppm for precursor and 20 ppm for fragment ions were required. A maximum of two missed cleavages was allowed for the tryptic digest. Cysteine carbamidomethylation was set as fixed modification, while N-terminal acetylation and methionine oxidation were set as variable modifications. MaxQuant processed output files can be found in Table S7, showing peptide and protein identification, accession numbers, % sequence coverage of the protein, q-values, and LFQ intensities. Contaminants, as well as proteins identified by site modification and proteins derived from the reversed part of the decoy database, were strictly excluded from further analysis.

### Metabolite Extraction and Profiling by Targeted LC-MS

About 30 mg of 10 unrelated and N_2_ shock frozen chRCC and corresponding healthy kidney tissues were used for metabolite profiling. Metabolite extraction and tandem LC-MS measurements were done as previously reported (Kurschner et al., 2017). In brief, methyl-tert-butyl ester (MTBE), methanol, ammonium acetate, and water were used for metabolite extraction. Subsequent separation was performed on an LC instrument (1290 series UHPLC; Agilent, Santa Clara, CA), online coupled to a triple quadrupole hybrid ion trap mass spectrometer QTrap 6500 (Sciex, Foster City, CA), as reported previously (Gielisch and Meierhofer, 2015). Transition settings for multiple reaction monitoring (MRM) are provided in Table S8.

The metabolite identification was based on three levels: (i) the correct retention time, (ii) up to three MRM’s and (iii) a matching MRM ion ratio of tuned pure metabolites as a reference (Gielisch and Meierhofer, 2015). Relative quantification was performed using MultiQuantTM software v.2.1.1 (Sciex, Foster City, CA). The integration setting was a peak splitting factor of 2 and all peaks were reviewed manually. Only the average peak area of the first transition was used for calculations. Normalization was done according to used amounts of tissues and subsequently by internal standards, as indicated in Table S8.

### Determination of Free and Total GSH in Plasma and Urine

Since the increased GSH level is a hallmark in renal oncocytoma and chRCC tissues, we asked, if this is also reflected in plasma and urine to establish a non-invasive metabolic marker. Therefore, plasma-(6 renal oncocytomas, 6 ccRCC, 12 pRCC, 6 chRCC, and 6 healthy) and urine specimens (8 renal oncocytomas, 20 ccRCC, 19 pRCC, 7 chRCC, and 20 healthy) were investigated. Free and total GSH were measured using a glutathione fluorescent detection kit (cat.no. EIAGSHF) according to the manufacturer’s protocol (Invitrogen, Carlsbad, CA).

### Enzyme Activities measurement

Sample preparation to spectrophotometrically assay enzyme activity was done as reported previously (Gielisch and Meierhofer, 2015). In brief, approximately 5 mg of tumor and healthy kidney tissues were homogenized and centrifuged at 600 g at 4 °C for 10 min. The protein concentrations of supernatants were further determined with a BCA assay (Thermo Fisher, Germany). For complex I, II, and V, 4 µg protein of each sample was used for the enzymatic activity measurement; for complex III, IV, and citrate synthase, 2 µg protein was used. Rotenone (10 µM), malonic acid (5 mM), antimycin A (5 µg/ml), potassium cyanide (500 µM), and oligomycin (5 µg/ml) served as specific inhibitors for complex I to V, respectively. The enzymatic activities of the hexosaminidases A and B were assayed as previously described by using 4-nitrophenyl N-acetyl-β-D-glucosaminide as an artificial substrate (Shibata and Yagi, 1996). Heat inactivation of hexosaminidase A was performed by pre-incubation of samples for 3 h at 48 °C (Grabowski et al., 1984).

### Whole Exome Sequencing and Mitochondrial Bioinformatics Analysis

DNA was isolated from the remaining pellets after metabolite extraction with a DNA purification kit according to the manufacturer’s protocol (QIAmp DNA Mini Kit for Tissues, QIAGEN, Hilden, Germany). In brief, the pellets were lysed by proteinase K and the RNA was removed by RNase, the RNA-free genomic DNA was then purified and eluted on QIAamp Mini spin columns for library preparation. The library preparation was performed according to Agilent’s SureSelect protocol (SureSelectXT Human All Exon V5, protocol version B4 August 2015) for Illumina paired-end sequencing, as reported previously (Kurschner et al., 2017). The fastq files were used as input for the MToolBox pipeline (Calabrese et al., 2014), in order to extract mitochondrial DNA sequences and quantify each variant allele heteroplasmy, as done previously (Kurschner et al., 2017). Copy number variations (CNVs) were inferred and visualized with the CNVkit (Talevich et al., 2016).

### Analysis of TCGA ChRCC RNA-seq Data

TCGA (ID: KICH) RNA-seq data were obtained from UCSC Xena (Goldman et al., 2018) (https://xenabrowser.net/). Accurate transcript quantification of chRCC (n=66) and controls (n=25) was based on the RNA-Seq by Expectation Maximization (RSEM) method (Li and Dewey, 2011).

### Experimental Design and Statistical Rationale, and Pathway Analyses

For proteome and metabolome data sets, a two-sample t-test was performed. Multiple test correction was done by Benjamini-Hochberg with an FDR of 0.05 by using Perseus (v1.6.0.2) (Tyanova et al., 2016). Significantly regulated proteins and metabolites were marked by a plus sign in the corresponding Tables S3 and S4. The Mann–Whitney *U* test was used to determine whether GSH levels from independent urine and plasma samples have the same distribution; the *p-value* significance cut-off was ≤ 0.01.

For comprehensive proteome data analyses, gene set enrichment analysis (GSEA, v3.0) (Subramanian et al., 2005) was applied in order to see, if *a priori* defined sets of proteins show statistically significant, concordant differences between chRCC and kidney tissues. Only proteins with valid values in at least seven of ten samples in at least one group with replacing missing values from the normal distribution for the other group were used (Table S3). GSEA default settings were applied, except that the minimum size exclusion was set to 10 and KEGG v6.2 was used as a gene set database. The cut-off for significantly regulated pathways was set to a p-value ≤ 0.01 and FDR ≤ 0.05.

### Data Availability

The datasets generated in the current study are available as supplementary files and in the following repositories:

WES files can be accessed via: https://www.ncbi.nlm.nih.gov/sra with the accession number: PRJNA413158.

Proteomics raw data have been deposited to the ProteomeXchange Consortium via the Pride partner repository (Vizcaino et al., 2013) with the dataset identifier PXD010391.

Metabolomics data have been deposited in the publically available repository PeptideAtlas with the identifier PASS01250 and can be downloaded via http://www.peptideatlas.org/PASS/PASS01250.

## Supplemental Information titles and legends

**Fig. S1. Volcano plot of transcript expression values between mutant (n=14, heteroplasmy ≥ 50%) and wt mtDNA (n=52) of a chRCC cohort, related to Figure 4.** Data were retrieved from TCGA. Shown are log_2_ expression values against the -log_10_ (*p*-value). Transcripts above the continuous line would be t-test significant (FDR ≤0.05).

**Fig. S2. Exome-based copy number variation analysis in chRCC, related to Figure 4.** chRCC specific monosomies of chromosomes 1, 2, 6, 10, 13, 17, and frequently of 21 were identified in all cases.

**Fig. S3. Glutathione levels in plasma and urine cannot be used as a marker to distinguish RCC’s from controls, related to Figure 5.** Urine samples were normalized to creatinine. *P*-values are: *P < 0.01.

**Fig. S4. Integration of protein- and metabolite abundances involved in amino acid metabolism. Simplified KEGG pathways show the (a) glycine and serine metabolism, (b) β-alanine metabolism, and (c) tryptophan metabolism, related to Figure 6.** * indicates significantly regulated proteins and metabolites; red = increased, blue decreased levels in chRCC, the value beside the metabolite indicate the fold change of the correspondent metabolite in chRCC versus kidney.

**Table S1. Clinical and pathologic features of the chRCC cohort, related to Methods.**

**Table S2. Protein ratios between chRCC and kidney tissues, related to Figure 1.** The significant threshold was set to *p*-value ≤0.05, FDR ≤0.05. Mitochondrial and non-mitochondrial proteins are indicated, the ratios between chRCC and kidney are shown for a high abundance in chromophobe RCC in red and a low abundance in chromophobe RCC in green.

**Table S3. Pathway enrichment analysis (GSEA) of all up-regulated (sheet 1) and down-regulated (sheet 2) pathways between chRCC and kidney tissues, related to Figure 1**. The significant threshold was set to *p*-value ≤0.05, FDR ≤0.05 and is highlighted in orange.

**Table S4. Identified metabolites in chRCC and kidney tissues, related to Figure 1.** Only valid values were used, shown as log_2_ transformed peak areas. The two-sample t-test was Benjamini-Hochberg corrected by applying an FDR of 0.05. The first transition, according to Supplemental Table S8 was used to calculate ratios and *p*-values. HMDB and KEGG identifiers are indicated.

**Table S5. The percentage of reconstructed genome covered by the assembly, the mean coverage depth, the number of contigs obtained and the best-predicted haplogroup are reported for each sample, related to Figure 4.** K = kidney, C = chRCC, same sample numbers are derived from the same patient. The next sheets show somatic and germline mtDNA mutations including the amino acid change, heteroplasmy rate, nucleotide variability and the disease score.

**Table S6. Proteome profiling and a subsequent GSEA analysis of UOK276 cells supplemented with 1% amino acids in the presence or absence of 2.5% BSA versus a complete medium, related to Figure 7.** The next two sheets show a pathway enrichment analysis (GSEA) of all up-regulated and down-regulated pathways of UOK276 cells in 1% amino acid with 2.5% BSA supplementation versus a complete medium. The significant threshold was set to p-value ≤0.05, FDR ≤0.05 and is highlighted in orange. The proteome profiles were done in triplicates.

**Table S7. MaxQuant output file featuring the proteome profile of chromophobe RCC’s and matching kidney tissues with LFQ intensities, related to Figure 1.**

**Table S8. Mass spectrometry settings for targeted metabolite profiling, related to Methods.** List of all metabolites with masses, MS conditions, and MRM ion ratios for the targeted LC/MS metabolite approach. Parameters for detection were optimized with prior measurements of the pure metabolites. If the pure substance was not available, parameters were created via database information (METLIN/HMDB), in-house solutions (MS/MS) or based on values of similar chemical structure (MRM’s not tuned in). For metabolites without a detectable third transition, a pseudo-MRM, measuring the precursor mass was used. A retention time (RT) of 0 min indicates that the metabolite was measured continuously, due to large peak widths or lack of optimized elution time information. Added internal standards are shown and the standard used for normalization within the method is highlighted as such. (DP = Declustering Potential, CE = Collision Energy, CXP = Collision Exit Potential).

