## Supplemental Information for "Metabolic reprogramming and elevation of glutathione in chromophobe renal cell carcinomas"

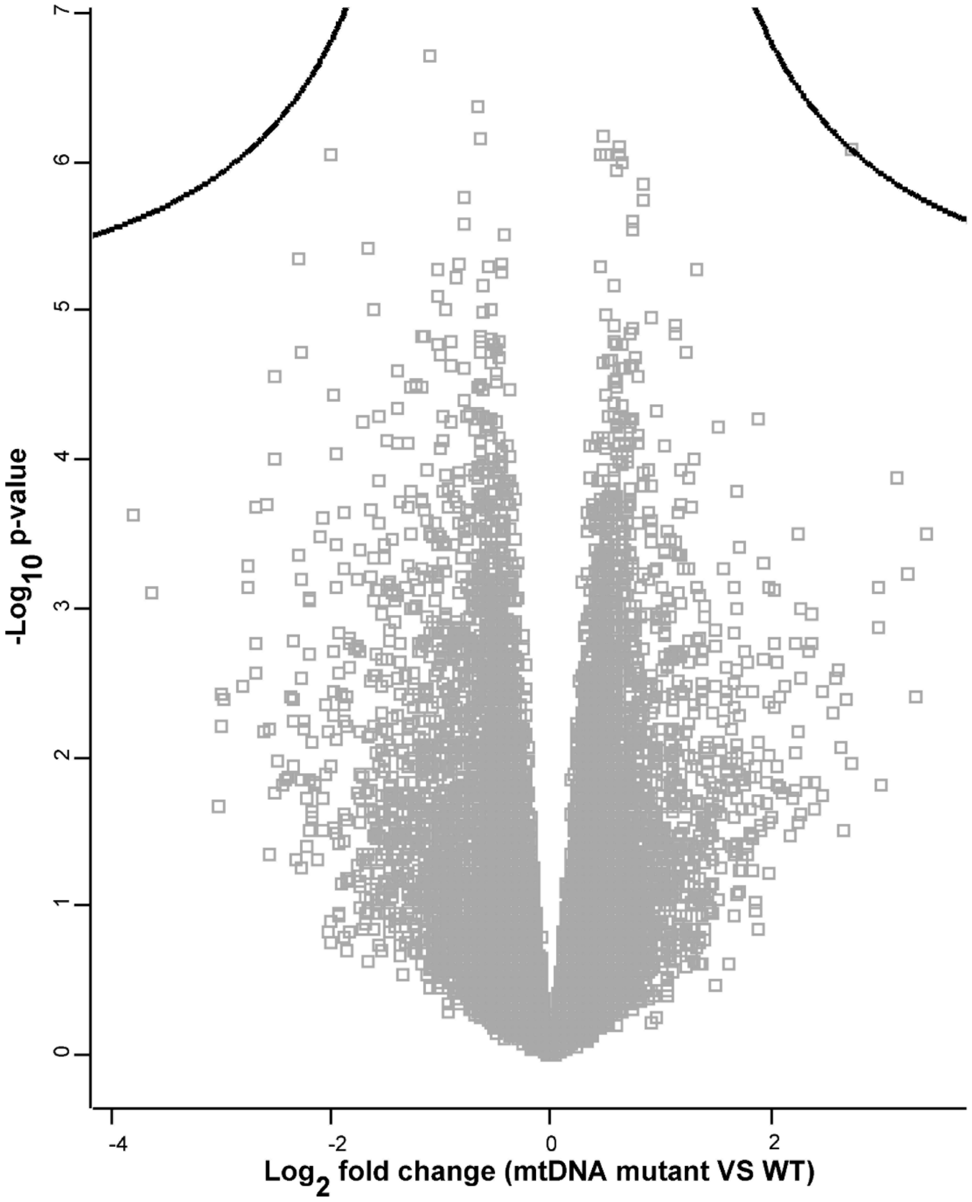


**Fig. S1. Volcano plot of transcript expression values between mutant (n=14, heteroplasmy ≥ 50%) and wt mtDNA (n=52) of a chRCC cohort, related to Figure 4.** Data were retrieved from TCGA. Shown are $\log_2$ expression values against the $-\log_{10}$ (*p*-value). Transcripts above the continuous line would be t-test significant (FDR ≤0.05).

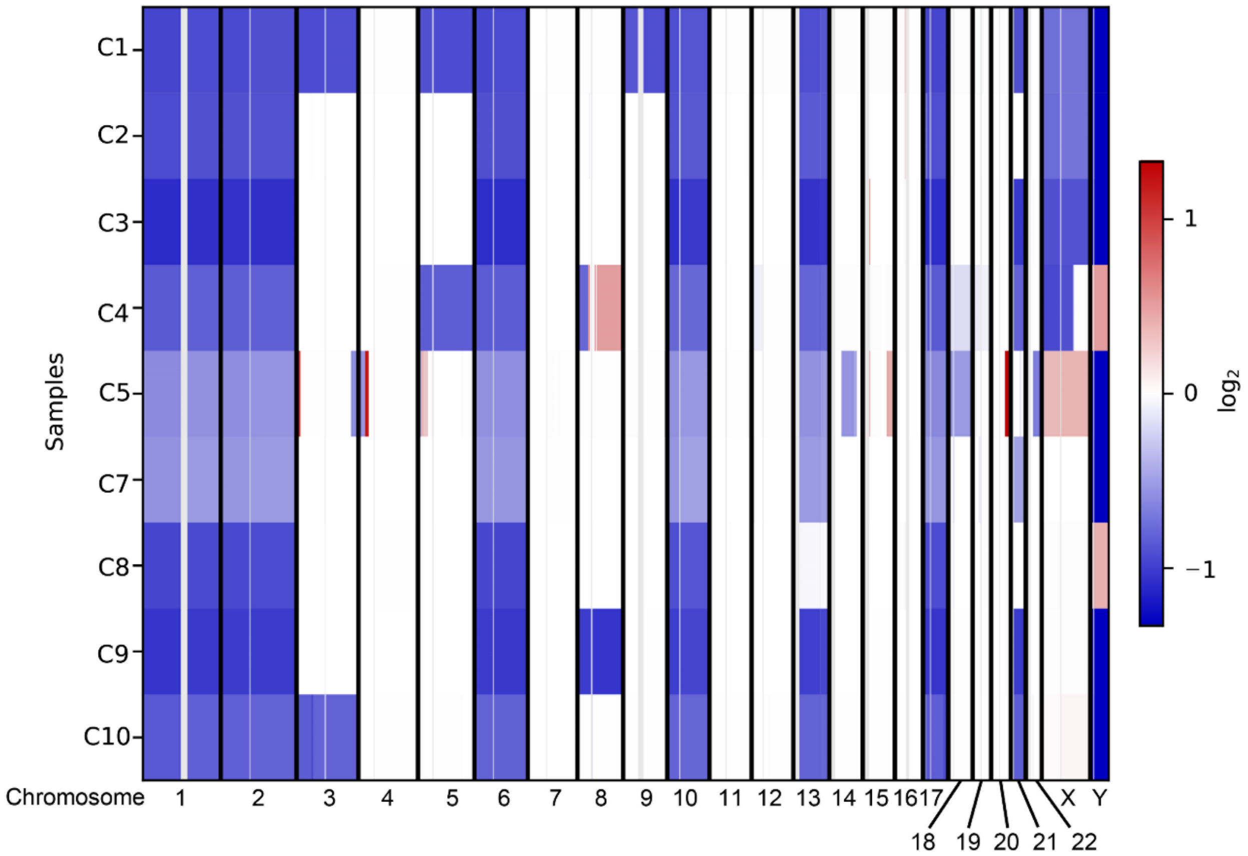


**Fig. S2. Exome-based copy number variation analysis in chRCC, related to Figure 4.** chRCC specific monosomies of chromosomes 1, 2, 6, 10, 13, 17, and frequently of 21 were identified in all cases.

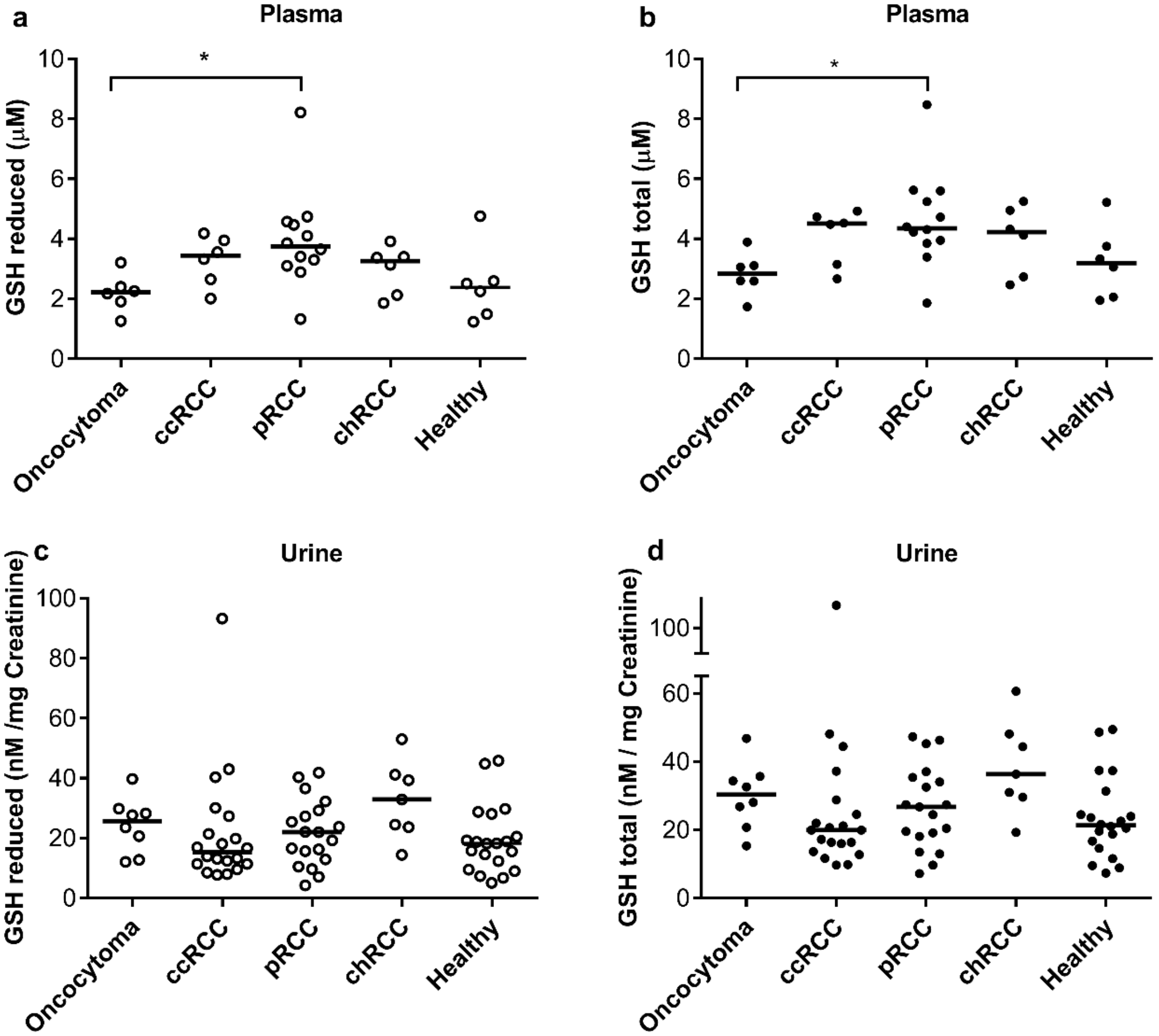


**Fig. S3. Glutathione levels in plasma and urine cannot be used as a marker to distinguish RCC's from controls, related to Figure 5.** Urine samples were normalized to creatinine. *P*-values are: *P < 0.01.

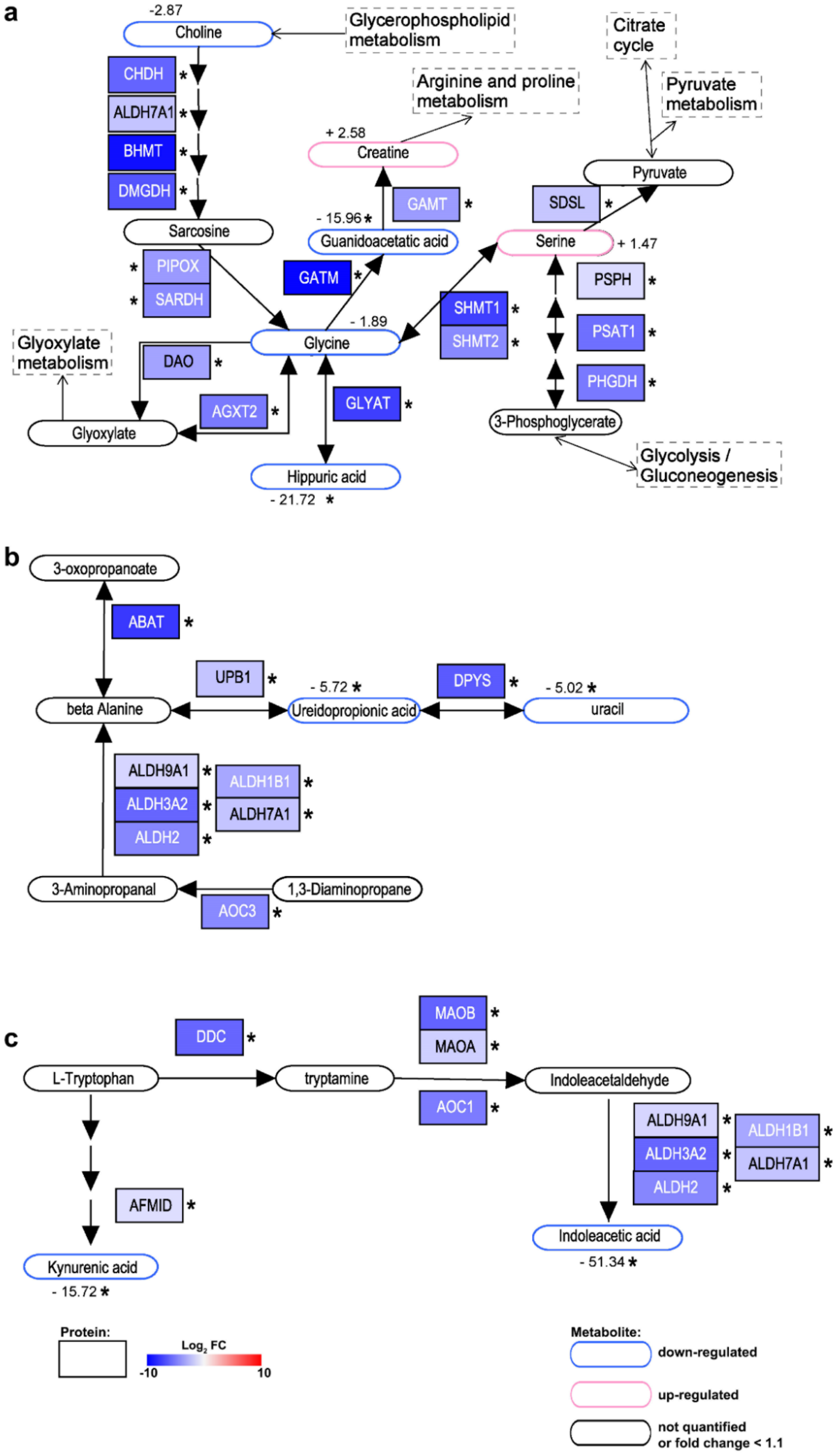

a
-2.87
Choline
Glycerophospholipid metabolism
CHDH
ALDH7A1
BHMT
DMGDH
Sarcosine
Arginine and proline metabolism
+ 2.58
Creatine
GAMT
- 15.96
Guanidoacetatic acid
Citrate cycle
Pyruvate metabolism
Pyruvate
SDSL
Serine
+ 1.47
PIPOX
SARDH
GATM
SHMT1
SHMT2
PSPH
PSAT1
PHGDH
- 1.89
Glycine
Glyoxylate metabolism
DAO
AGXT2
GLYAT
Glyoxylate
3-Phosphoglycerate
Glycolysis / Gluconeogenesis
Hippuric acid
- 21.72
b
3-oxopropanoate
ABAT
UPB1
- 5.72
DPYS
- 5.02
beta Alanine
Ureidopropionic acid
uracil
ALDH9A1
ALDH3A2
ALDH2
ALDH1B1
ALDH7A1
3-Aminopropanal
1,3-Diaminopropane
AOC3
c
DDC
MAOB
MAOA
L-Tryptophan
tryptamine
Indoleacetaldehyde
AOC1
ALDH9A1
ALDH3A2
ALDH2
ALDH1B1
ALDH7A1
AFMID
Indoleacetic acid
- 51.34
Kynurenic acid
- 15.72
Protein:
Log2 FC
-10
10
Metabolite:
down-regulated
up-regulated
not quantified or fold change < 1.1

**Fig. S4. Integration of protein- and metabolite abundances involved in amino acid metabolism. Simplified KEGG pathways show the (a) glycine and serine metabolism, (b) β-alanine metabolism, and (c) tryptophan metabolism, related to Figure 6.** * indicates significantly regulated proteins and metabolites; red = increased, blue decreased levels in chRCC, the value beside the metabolite indicate the fold change of the correspondent metabolite in chRCC versus kidney.

**Table S1. Clinical and pathologic features of the chRCC cohort, related to Methods.**

| Case ID | Age at surgery | Gender | Pathologic T | Grade | Tumor Size (mm) | Nephrectomy |
|---|---|---|---|---|---|---|
| C1 | 49 | Female | pT1a | 2 | 28 | radical, left |
| C2 | 37 | Female | pT2a | 2 | 75 | radical, right |
| C3 | 42 | Female | pT2a | 2 | 98 | radical, left |
| C4 | 50 | Male | pT3a | 2 | 100 | radical, right |
| C5 | 71 | Female | pT3a | 3 | 70 | radical, right |
| C7 | 38 | Female | pT1b | 1 | 60 | partial, right |
| C8 | 58 | Male | pT1b | 1 | 65 | radical, left |
| C9 | 36 | Female | pT2b | 2 | 110 | radical, left |
| C10 | 46 | Male | pT2a | 0 | 74 | radical, right |

**Table S2. Protein ratios between chRCC and kidney tissues, related to Figure 1.** The significant threshold was set to *p*-value ≤0.05, FDR ≤0.05. Mitochondrial and non-mitochondrial proteins are indicated, the ratios between chRCC and kidney are shown for a high abundance in chromophobe RCC in red and a low abundance in chromophobe RCC in green.
